# Undergraduate student practicals generate high-quality data for microbiome research

**DOI:** 10.64898/2026.01.20.700693

**Authors:** Isabella Wilson, Tahlia Perry, Frank Grutzner

## Abstract

The increasing prominence and accessibility of microbiomics has provided an opportunity for authentic research experiences in the undergraduate practical classroom. In recent years, this approach has contributed to published research projects. However, there is little information evaluating the quality of student-generated data compared to that of trained researchers. To investigate this, we designed an undergraduate practical component in which 37 final-year genetics students generated microbial profiles of 22 echidna scats using matched samples that were also profiled by an experienced researcher. DNA yield, 16S PCR success, sequencing library size and microbial diversity were compared between the groups in order to assess both the ability and accuracy of students in characterising faecal microbiota. Our research revealed that students were able to produce research-grade microbiome data comparable to a postgraduate researcher. Importantly, we found that students did not introduce contamination at a higher rate than the trained researcher. These findings reinforce that the undergraduate classroom is a valuable approach for microbiome research in addition to its benefits for student engagement and experience. The design and successful implementation of these practicals provide a template for a variety of research-led microbiome teaching.

## Introduction

Over the past 30 years, the close relationship between microbiomes and health has been revealed through an explosion of research, gaining recognition from the scientific community and the general public. The importance of microbiota extends beyond humans with advances also being made in the area of animal microbiomes. Microbiota in various animal species have been associated with resilience to pathogens, reproductive success, and social behaviours such as communication^1–3^. Animal microbiome studies provide information that is being used to understand and improve the health of species ranging from livestock to endangered wildlife^4^.

This increasing recognition of the microbiome, as well improved affordability of tools needed for microbiome research such as DNA extraction kits and next-generation sequencing technologies^5^, has resulted in many universities introducing microbiomics into undergraduate curricula. A innovative way in which this has been achieved is by incorporating authentic research experiences into practical classes, for example through course-based undergraduate research experiences (CUREs) or inquiry-based learning^6^. These approaches to practical design allows students to be participants in the scientific process and can involve generating new data to address unanswered research questions^7^. Compared with traditional “cookbook” practicals, CUREs have been shown to improve grades in companion courses, reduce barriers to research participation for disadvantaged student groups, and increase student retention rates^8–10^.

There is a large diversity of microbiome-based CUREs documented in the literature with programs covering oral, faecal, genital, animal, soil, water, plant, and food microbiomes. These CUREs include any or all of the steps typical of microbiome research: sample collection (via self-collection, field samples, or use of existing samples obtained through ongoing research projects); bacterial culture/microscopy; DNA extraction; library preparation/16S PCR; sequencing; and data analysis^11^. Although authentic research experiences have documented benefits for student learning, the validity of the scientific data produced by microbiome-based CUREs has not been evaluated. Microbiome research presents a range of difficulties surrounding contamination which may be compounded by the student practical setting. It is unknown whether students are more prone to introducing contaminating microbes from their skin, hair, or breath as they are still in the process of learning optimal laboratory technique. Additionally, the student laboratory environment itself may contain a higher incidence of contaminating bacteria due to other practicals occurring in the shared teaching space. It is important to understand the impact of these potential contamination sources, particularly for data that contribute to existing research projects and are included in peer reviewed journal articles^12–16^.

To measure the quality of microbiome data generated in a professional versus student laboratory environment, we designed a practical unit in which undergraduate students extracted and sequenced bacteria from short-beaked echidna scats. Scat samples were sourced through the citizen science project EchidnaCSI^17^. This project has facilitated the collection of echidna scats from across Australia by the general public and has already provided new insights into echidna biology and diversity^18^; however, the large volume of samples acquired through EchidnaCSI and other similar citizen science initiatives requires innovative approaches for high-throughput sample processing and sequencing. A set of twenty-two echidna scat samples were extracted by undergraduate genetics students as well as a postgraduate researcher, after which they were sequenced using an Oxford Nanopore Technologies MinION™. The ability of the students to complete each part of the experimental workflow was compared to the trained researcher. The diversity and taxonomic composition of the student and researcher samples were evaluated.

## Methods

### Study design

Twenty-two echidna scats were collected in the Flinders Ranges, South Australia, through the EchidnaCSI citizen science project before being stored at -80°C. Each sample was divided so that they could be processed by one researcher and at least two undergraduate student groups. This allowed for researcher-student as well as student-student comparison and increased the likelihood of at least one student-processed sample being successfully sequenced. “Researcher” samples were extracted by a postgraduate researcher with experience in echidna scat microbiomics. “Student” samples were extracted as part of the practical component of the final year undergraduate Genetics course at the University of Adelaide. For this study, we included data from the 2023/24 student cohort; 22 students participated in 2023 and 15 students participated in 2024. However, the practical component is ongoing and students continue to generate new data.

### Experimental methods

For the researcher-extracted samples, DNA extractions were performed in a PC2-certified laboratory at the University of Adelaide. The laboratory bench and equipment were cleaned with 10% bleach prior to use except for pipettes which were cleaned with 70% ethanol. For the student-extracted samples, extractions took place within a PC1-certified undergraduate teaching laboratory. Students were instructed to clean their benches and equipment with 70% ethanol due to the unavailability of bleach as a cleaning agent in the undergraduate laboratory setting.

Total genomic DNA was extracted from scat samples using the QIAGEN QIAamp Fast DNA Stool Mini Kit according to the manufacturer’s instructions, except as outlined below. For all samples, scats were homogenised using a mortar and pestle nestled in an esky of dry ice prior to the addition of 1.4 mL InhibitEX buffer. Samples were centrifuged for 3 min at maximum speed. 800 µL supernatant was transferred to a clean 1.5 mL tube before being centrifuged for 3 min at maximum speed. 600 µL supernatant was transferred into a clean 2 mL tube along with Proteinase K. After elution, whole DNA was run on a 1.5% gel for 30 min at 100 V. Concentration was measured using a Qubit fluorometer before being stored at -20°C (quantification step performed by researcher for all samples). The researcher and each student group prepared an extraction blank control in order to capture laboratory-based contamination. For these samples, no echidna scat was added to the lysis tube at the beginning of the extraction protocol.

Library preparation approaches differed slightly between samples sequenced in 2023 and 2024 due to changes to Nanopore kits within this period. Information regarding library preparation and sequencing kits for each sample can be found in the supplementary metadata file. Library preparation for the 2023 cohort was performed using the Nanopore 16S Barcoding Kit 1-24 (SQK-16S024) whereas in 2024 the Nanopore 16S Barcoding Kit 24 V14 (SQK-16S114.24) was used. For researcher samples, the researcher performed all library preparation steps. The changing roles of the students and the researcher in preparation of the student samples are summarised in Table 1.

**Table 1:** Different roles of students and researcher in library preparation as a result of changes to Nanopore protocol.

| Step | 2023 | 2024 |
| --- | --- | --- |
| 1 | 16S PCR; Researcher | 16S PCR; Researcher |
| 2 | 16S gel; Researcher | 16S gel; Researcher |
| 3 | Clean-up; Student | Quantification; Student |
| 4 | Quantification; Student | Pooling; Researcher |
| 5 | Pooling; Researcher | Clean-up; Researcher |
| 6 | Sequencing; Researcher | Sequencing; Researcher |

For all samples, sequencing was performed on a Nanopore MinION Mk1C. Samples from 2023 were sequenced using R9.4.1 flow cells whereas samples from 2024 were sequencing using R10.4.1 flow cells.

### Biosafety considerations

The practical series was targeted at final year undergraduate Genetics students who had previously received considerable safety training and were familiar with the fundamentals of safe behaviour in laboratory settings. Students wore necessary protective personal equipment (PPE) including gloves, safety glasses, protective footwear, and laboratory coats. Laboratory benches and equipment were decontaminated with 70% ethanol before and after each session. Students were instructed to wash their hands before leaving the teaching laboratory. Phones and other personal devices were prohibited to prevent surface contamination. Biological waste was handled and disposed of in accordance with established University of Adelaide teaching laboratory protocols.

### Data processing and statistical analysis

Basecalling and barcode demultiplexing was performed using MinKNOW™. Demultiplexed reads were processed using MetONTIIME, a pipeline that adapts QIIME2 for Nanopore 16S data^19,20^. Filtering was performed in order to retain reads between 1000-1800 bp and with a minimum quality-score of 10. *De novo* clustering was performed at 95% identity. Taxonomic classification was performed using the SILVA 138 SSURef NR99 full-length 16S reference database^21^. Diversity analysis was performed at a depth of 1800 reads at the genus level. Alpha diversity was estimated using observed operational taxonomic units (OTUs), Shannon diversity, and Pielou’s evenness; beta diversity was estimated using Bray-Curtis dissimilarity and Jaccard dissimilarity.

Further statistical analysis and data visualisation was performed using R. For analysis of DNA concentration, average concentration was calculated per scat sample for the students and the researcher in order to account for repeated measurements in student group. These sample means were assessed for significance using a paired Wilcoxon signed-rank test. To analyse alpha diversity, student replicates were matched to researcher replicates according to scat ID and the difference between alpha diversity values was calculated. A Wilcoxon signed-rank test was used to determine whether the median of these differences was significantly different to zero. For beta diversity, principal coordinate analysis (PCoA) and hierarchical clustering were used to determine the relative similarity between replicates of each sample. Alpha and beta diversity were visualised using ggplot2^22^ and ggdendro^23^.

For taxonomic analysis, bar plots showing the relative abundances of genera within experimental samples and total abundances of genera within extraction blank controls were generated using phyloseq^24^ and microViz^25^. phylosmith^26^ was used to identify taxa unique to student or researcher replicates. In order to identify differentially abundant genera, the feature table was glommed to the genus level; absolute counts were transformed into relative counts; low-abundance taxa (mean relative abundance < 0.001) were filtered out; and pseudo-counts were added. The relative abundance of each genus within each scat was calculated for the researcher and student replicates, and the average abundance of each genus was taken for student replicates. From this, log ratio differences in genus abundance between student and researcher replicates were calculated. Log ratios were aggregated by genus across all samples, allowing for assessment of overall student-researcher differences (rather than within-sample differences). To test these for significance, a Wilcoxon signed-rank test was used to determine whether the median log-ratio for each genus was significantly different to zero. The resulting p-values were adjusted for multiple testing using the Benjamini-Hochberg procedure. decontam^27^ was used to identify potential contaminant taxa based on the contents of the extraction blank controls and PCR negative (no template) controls; the prevalence method was used with a threshold of 0.5.

Code for all statistical analysis and data visualisation can be found at the following GitHub repository: https://github.com/isa-wilson/microbiome_pracs/.

## Results

Two four-hour practical sessions were conducted as part of the University of Adelaide 3^rd^-year Genetics course over two consecutive years (2023 and 2024) (Fig. 1). Practical manuals for each session can be found in the supplementary data. A total of 96 replicates originating from 22 short-beaked echidna scat samples were processed by 37 students and one postgraduate researcher. Over 4.5 million reads were generated across five Nanopore sequencing runs.

**Figure 1:**
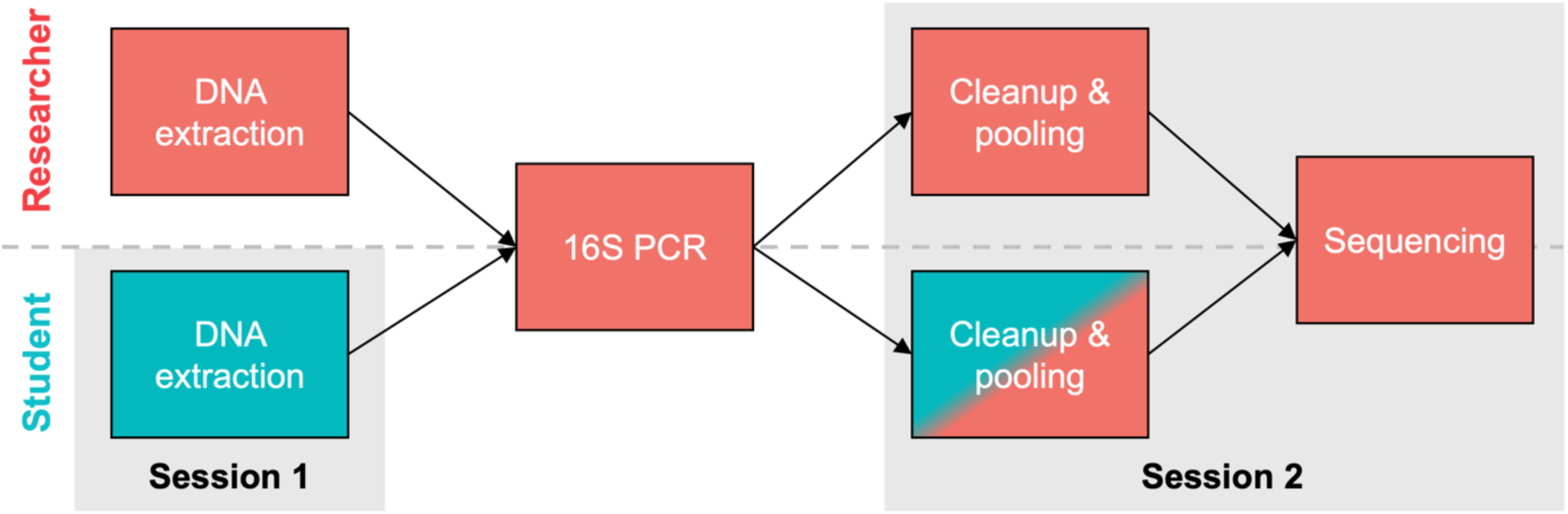
Schematic summarising the main steps of the experiment. Tasks performed by students are highlighted in blue. Tasks performed by the researcher are highlighted in red. Steps taking place within practical sessions are outlined in grey.

### Undergraduate students generate research-grade data

We first examined the students’ ability to successfully complete each main step of the typical 16S workflow: DNA extraction, polymerase chain reaction (PCR), and sequencing.

DNA extraction success was quantified using DNA yield, which refers to the quantity of whole DNA (including from bacteria, echidna, and other sources) that was isolated from the scat samples (Fig 2a). The DNA concentration of researcher-extracted samples (median 6.22 ng/μL) was significantly higher than that of students (median 3.43 ng/μL) (p = 0.039; Wilcoxon signed-rank test).

**Figure 2:**
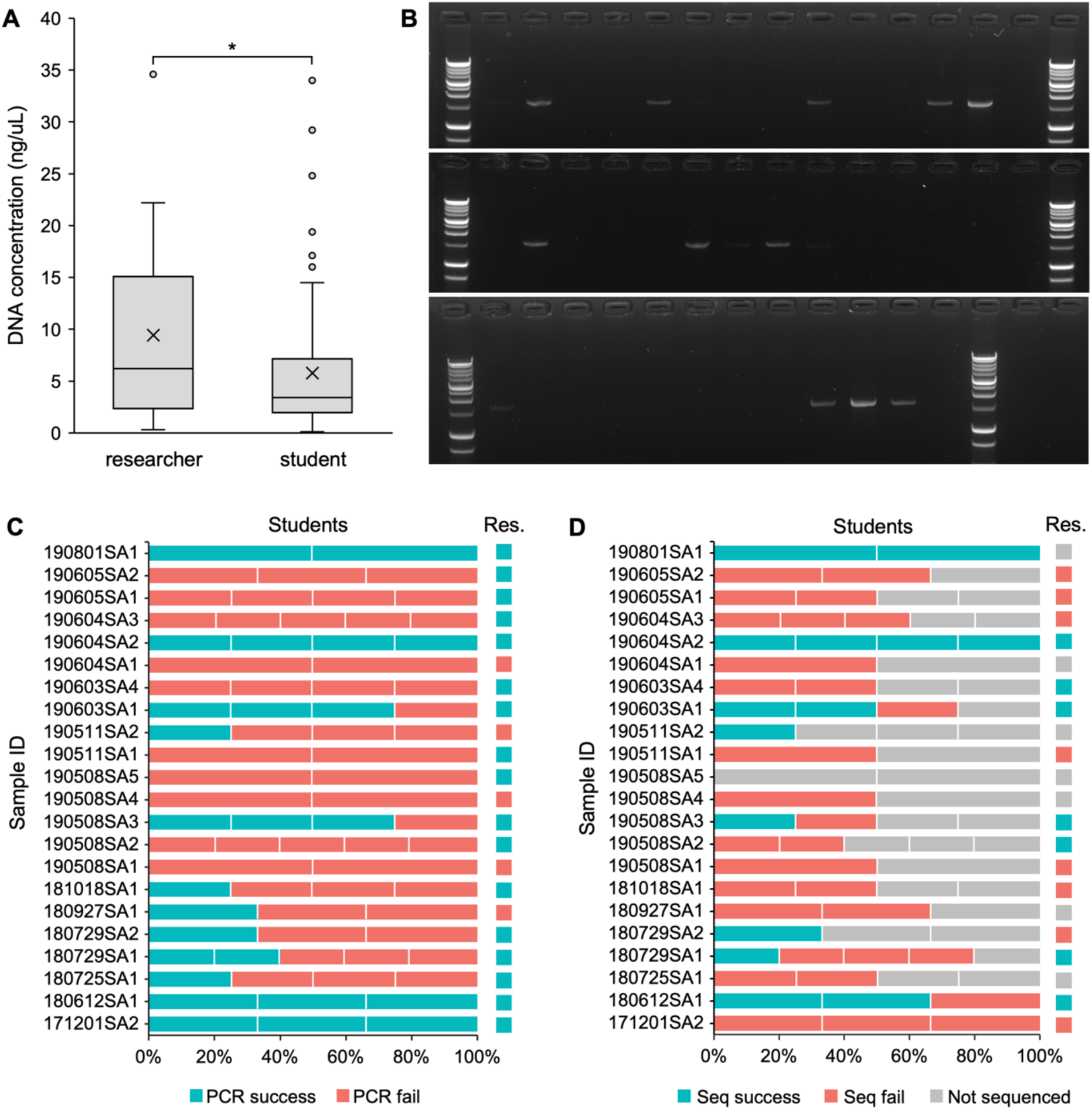
Comparison of experimental efficiency between students and researcher for each step of the microbiome workflow. **A)** Box-and-whisker plot comparing DNA concentration of extractions. Cross indicates mean, asterisk indicates statistical significance (p-value < 0.05). **B)** Gel electrophoresis showing 16S PCR products from student extractions. A band approximately 1500 bp in size can be seen for some samples. **C)** Bar plot showing per-sample PCR success rates for students vs researcher (abbr. “Res.). **D)** Bar plot showing per-sample sequencing success rates for students vs researcher.

The 16S PCR is a critical step in the microbiome workflow as it generates amplicons for sequencing. A successful PCR can be measured by gel electrophoresis (Fig 2b). The presence of a band of the expected size (approximately 1500 bp) demonstrates that intact bacterial DNA was successfully extracted from the scat samples and that contamination by PCR inhibitors was minimal. PCRs using researcher-extracted DNA achieved a success rate of 77.27%, whereas student-extracted DNA averaged only 33.78% success (Fig 2c). For the majority of scats, the student groups produced uniform results (i.e. all succeeded or all failed). At least one student DNA extract was PCR-amplified for 12 out of the 22 scat samples (54.44%). Considering that the student and researcher extractions yielded significantly different DNA concentrations, we tested whether there was a relationship between DNA concentration and PCR success for student replicates; however, we did not find evidence to support this (p = 0.091; Wilcoxon rank-sum test).

Not all samples could be sequenced due to limited space on the flow cell. As such, we prioritised sequencing at least one replicate for each scat sample. Additionally, replicates with a very low concentration (<0.5 ng/μL) after the cleanup stage were generally not sequenced. In total, 14 researcher and 24 student samples were sequenced (Fig 2d). We defined sequencing to be successful if a sampling depth of ≥ 1800 reads was achieved. This was selected based on alpha rarefaction plotting; this depth was where alpha diversity metrics (observed OTUs, Shannon diversity) began to plateau (Supplementary Fig. S1) while retaining enough OTUs to allow for meaningful statistical analysis of bacterial diversity. In order to compare sequencing success across students and researchers, we restricted our analysis to samples that had successfully undergone 16S PCR amplification as these could be expected to contain the DNA necessary for sequencing. Of the 14 samples extracted by the researcher that were successfully PCR amplified, 7 achieved a sampling depth of ≥ 1800 (50%). Interestingly, this was elevated for the students; of the 24 PCR-success samples that were sequenced, 14 achieved a sampling depth of ≥ 1800 (58.33%). This suggests that so long as a sample is able to be successfully PCR-amplified, researchers and students have a similar chance of achieving sequencing success.

### Samples isolated by students and a trained researcher show compositional similarity

To investigate bacterial community structure, we compared the student and researcher-isolated microbiomes using beta diversity and taxonomic analyses. Beta diversity analysis, in this study using Bray-Curtis dissimilarity and the Jaccard index, quantifies dissimilarity between sample groups and provides insight into compositional differences. For both Bray-Curtis and Jaccard dissimilarity, PCoA plotting showed that replicates deriving from the same scat clustered together (Fig. 3), indicating that for a given scat sample, researcher and student replicates were more similar to each other than to any other scat sample. This was confirmed with hierarchical clustering (Supplementary Fig. S2). Some samples were more tightly clustered than others; for example, all five 190604SA2 replicates were very tightly clustered whereas the 190801SA1 replicates were more dissimilar.

**Figure 3:**
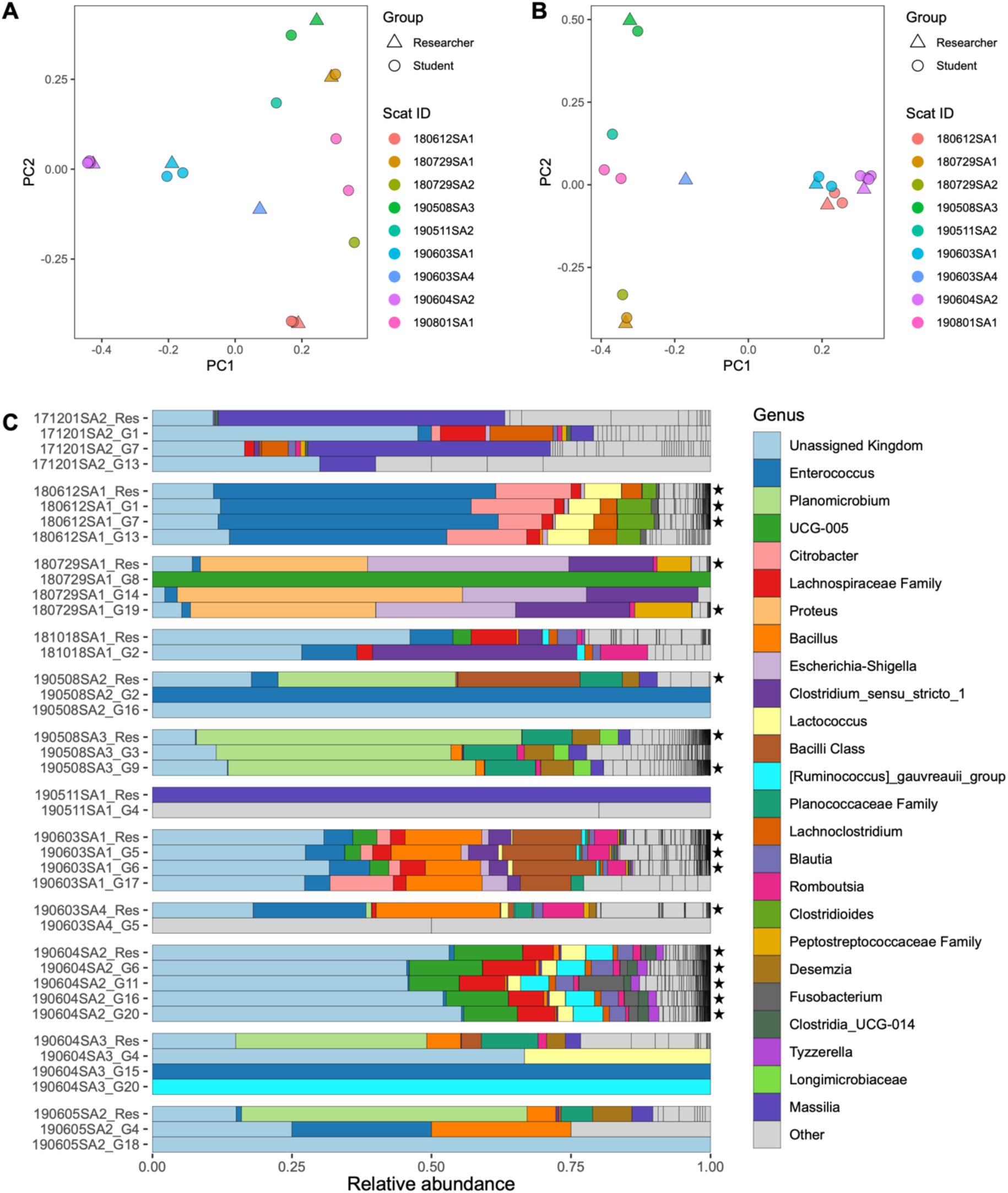
Compositional analysis of student and researcher-isolated samples. **A)** PCoA plot showing Bray-Curtis distances between samples. **B)** PCoA plot showing Jaccard distances between samples. Beta diversity metrics were calculated at the genus level using a sampling depth of 1800. **C)** Relative abundance of genera present within student and researcher replicates. Replicates are grouped according to scat sample. Top 20 genera are displayed to the right of the plot in order of abundance. Star indicates samples that were included in diversity analysis, i.e. sampling depth ≥ 1800.

We next compared the taxonomic composition of the student and researcher-derived samples. As expected from our beta diversity analysis, student and researcher replicates from the same scat sample were overall compositionally similar (Fig. 3c). The exception to this was for replicates with a low sequencing depth; for these samples, only a few genera were captured. Interestingly, there was a large proportion of features that could not be assigned to a Kingdom; for some samples (e.g. 190604SA2), these unassigned features comprised >50% of all observed taxa. In order to detect significant differences in the abundance of particular genera between student and researcher replicates, we calculated log ratio differences for the abundance of each genus in the student and researcher replicates of each scat. We then tested whether these log ratio differences were significantly different to zero; however, we observed no significant differences in abundance for any genera.

### Number and distribution of bacterial genera differ between student and researcher-generated data

Another aspect of microbiome community structure is alpha diversity, which estimates richness and evenness within samples. We examined the alpha diversity of student and researcher-isolated samples using observed operational taxonomic units (OTUs), which calculates the number of distinct taxa within each sample (richness); Pielou’s evenness, which measures whether the abundance of each taxon is evenly distributed; and Shannon diversity, which measures both richness and evenness (Fig. 4). For all metrics, diversity values were generally higher for student replicates compared to researcher replicates. Uneven sampling as a result of multiple replicates for the student group and only one replicate for the samples isolated by the researcher meant that the alpha diversity scores could not be directly compared. Instead, the difference between student and researcher replicates was calculated for each scat, and a Wilcoxon signed-rank test was used identify systematic differences between researcher and student-derived replicates. We found significant differences in observed OTUs (p = 0.032) and Shannon diversity (p = 0.0098), but not Pielou’s evenness (p = 0.160). These results suggest that while student and researcher-derived replicates for each scat samples are compositionally similar, significant differences are present in terms of the number of taxa captured as well as the distribution of abundances across the taxa.

**Figure 4:**
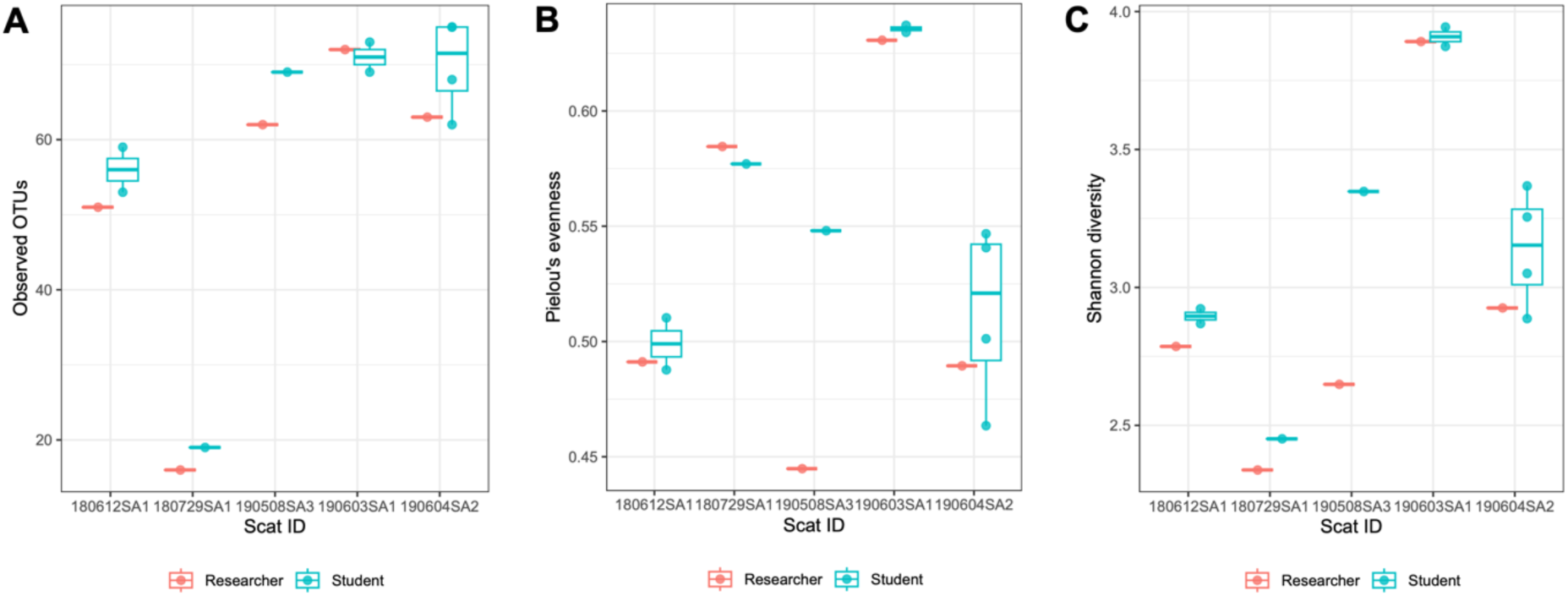
Alpha diversity analysis of student and researcher-isolated samples. **A)** Boxplot comparing per-sample observed OTUs between researcher and student replicates. **B)** Boxplot comparing per-sample Pielou’s evenness between researcher and student replicates. **C)** Boxplot comparing per-sample Shannon diversity between researcher and student replicates. All metrics were calculated at the genus level using a sampling depth of 1800.

### Samples isolated by students do not contain higher levels of contamination than researcher

The ubiquity of microbial DNA makes contamination a persistent issue in microbiome research. In order to gain a better understanding of potential contamination, we calculated the proportion of genera that were only observed in either student or researcher replicates. For researcher samples, 0.25% of total abundance was comprised by researcher-specific taxa whereas 99.75% was also present in student samples. For student samples, 0.33% of total abundance was comprised by student-specific taxa whereas 99.67% was also present in researcher samples. The similarity of these values suggests that students were not contributing environmental contaminants at a higher rate than the researcher. Next, we compared extraction blank controls and experimental samples. The concentration of all extraction blank controls was lower than the Qubit detection range (<0.05 ng/μL) which resulted in low read depth and in some cases a complete failure to undergo sequencing. Of the 7 extraction blank controls that were able to be sequenced, feature abundances ranged from 1 – 21 (Fig. 5). Only 25 genera were detected in extraction blank controls, all of which were also present in experimental samples except for *Aggregatibacter*, *Veillonella*, *Haemophilus*, and *Neisseria*. We used the R package decontam to identify contaminants by assessing whether any taxa were more prevalent in extraction blank controls and PCR negative controls than experimental samples; however, none could be identified. We also checked for differential abundance of particular taxa that could represent contamination, such as chloroplast and taxa that could not be assigned to a kingdom; again, no significant differential abundance was detected. Overall, we did not find evidence that students contributed increased levels of contamination.

**Figure 5:**
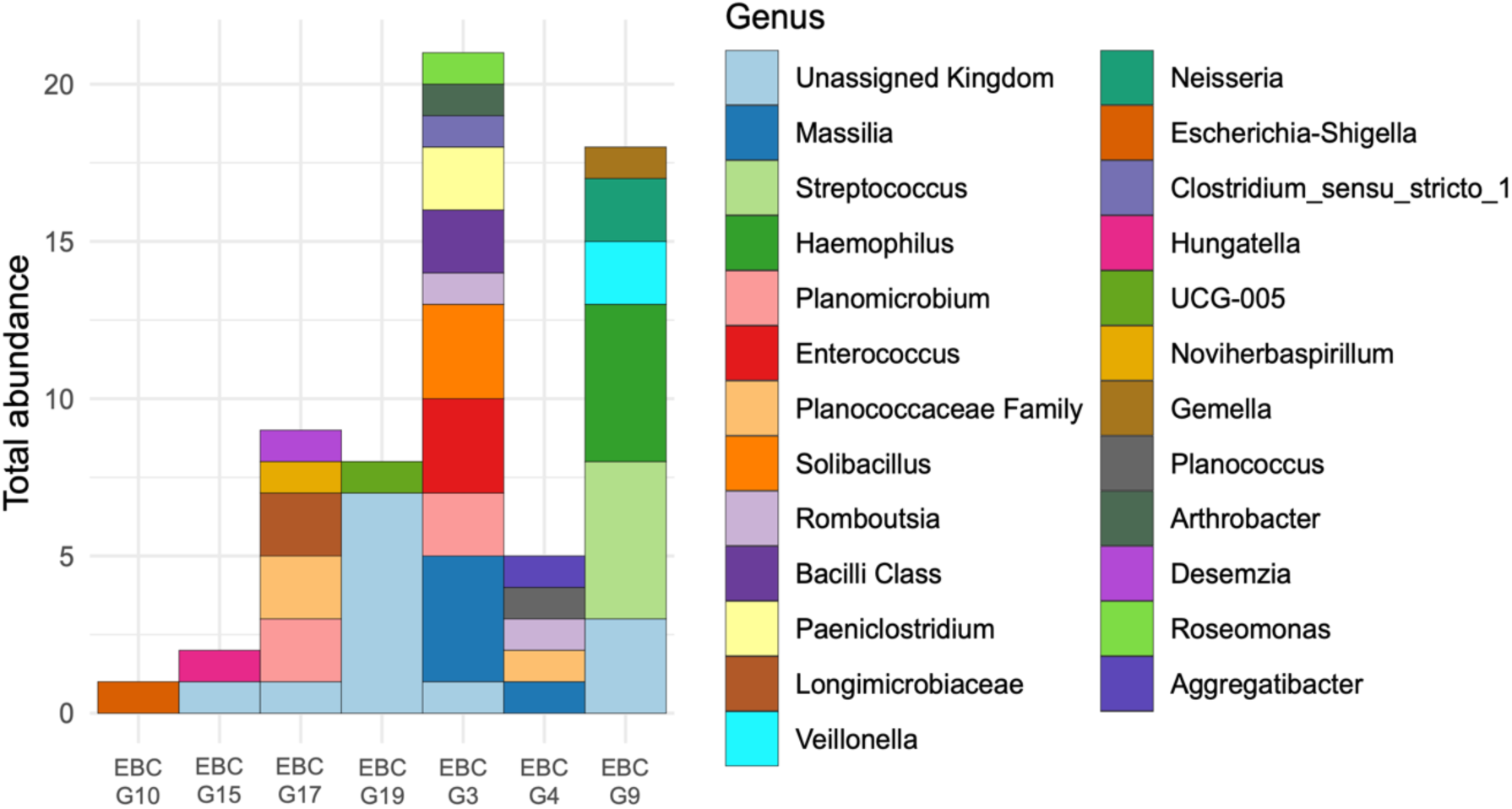
Total abundance of genera present within student extraction blank controls. Genera are displayed to the right of the plot in order of abundance.

## Discussion

Microbiome analysis of environmental samples has increased dramatically over the past decade and has clearly demonstrated the importance of the microbiome for both health and the environment. The streamlining of 16S rRNA workflows, cheaper sequencing and user-friendly analysis software for metabarcoding samples makes it feasible to use classroom-based practicals to generate microbiome research data. In addition to advancing research, these authentic research experiences also provide a more engaging learning environment for undergraduate students. Here we compared metabarcoding data generated by students and researchers from echidna scat samples. Our results highlight the potential and limitations of classroom-based research and show that a well-designed workflow can enable the generation of novel, high-quality research data.

The ability of students to execute the 16S workflow to the same standard as the researcher varied between the different experimental steps (Fig. 2). Student extractions produced significantly less DNA than the researcher, presumably due to a lack of experience with sample homogenisation technique (for example, the bacterial cells may not have been broken down sufficiently to allow for the isolation of DNA). Protocols using automated extraction devices (for example using a tissue homogenizer) could therefore potentially mitigate this issue. The 16S PCR stage proved to be the most challenging for the student groups. PCR results were expected to be variable for both researcher and student groups due to the inherent difficulty of working with echidna scat samples; faecal samples are notorious for containing PCR inhibitors, and DNA extraction kits are typically optimised for human stool samples rather than animal scats^28,29^. The PCR success rate for student samples (33.78%) was less than half of that of the researcher (77.27%). We did not find a correlation between the low DNA concentration in the student-extracted samples and PCR success rate, suggesting issues of DNA quality rather than DNA quantity. Previous microbiome-based CUREs have highlighted PCR as a step that is frequently challenging for students, to the extent that spare sessions are scheduled to allow students to repeat experiments^14,30^. In light of this, the low PCR success rate observed in this study may not be unusual.

Surprisingly, for two samples, a student group succeeded where the researcher failed, illustrating the unpredictable nature of processing environmental samples. Once students progressed past the PCR stage, success rates improved, with students and researchers exhibiting similar sequencing success. This suggests that PCR is a “hurdle” step in the protocol for classroom-based research; once 16S amplicons are successfully generated, researchers and students can achieve similar success. Our results support the use of at least 3 replicates and/or flexible timetabling for repeat experiments in order to facilitate a level of sequencing success on par with a professional researcher. It should also be noted that the number of students in a given practical cohort allows for the parallel processing of several replicates per scat sample, potentially improving throughput in comparison to a single researcher despite lower experimental efficiency.

Discrepancies between student and researcher replicates were also observed with respect to bacterial alpha diversity. Microbial richness and Shannon diversity were significantly different between the two groups, with students generally producing elevated values for both metrics. This meant that samples generated by students captured more of the bacterial genera within the scat sample. The reasons for the discrepancy in alpha diversity remain unclear, though the DNA extraction process may play a role; research has demonstrated that out of a range of experimental factors, DNA extraction has a comparatively large influence on microbiome profiling^31–33^. Considering that the extraction stage of the protocol had the least researcher involvement (as compared to PCR or sequencing), it is plausible that differences in extraction technique between the students and the researcher resulted in elevated bacterial richness.

Despite the differences in experimental efficiency and alpha diversity, compositional analysis including beta diversity and taxonomy demonstrate that students were able to generate similar microbial profiles to the researcher using the same scat sample. This was evidenced by PCoA and hierarchical clustering, which confirmed that replicates from the same scat were more similar to each other than to any other sample, and that no genera were identified as significantly differentially abundant between researcher and student replicates (Figure 3). Furthermore, the taxonomic makeup of the replicates was generally in line with previous studies characterising echidna gut microbiota^18,34^. The top 10 most abundant genera across all samples were *Enterococcus*, *Planomicrobium*, *UCG-005*, *Citrobacter*, an unknown *Lachnospiraceae* sp., *Proteus*, *Bacillus*, *Eschericia-Shigella*, *Clostridium_sensu_stricto_1*, and *Lactococcus*, all of which have been identified as members of the echidna gut microbiome in previous studies and are documented commensals of the gut microbiota of humans and/or animals^35–43^. Some genera such as *Citrobacter* were observed in higher abundances compared with previous studies. This genus is a common inhabitant of soil and gut microbiota, suggesting that its presence is not necessarily due to contamination^44^. While the reason underlying this elevated abundance is unknown, it may relate to the use of an updated taxonomic classifier; alternatively, it may reflect the inherent differences between the Nanopore approach of full-length 16S gene sequencing versus the Illumina approach of sequencing specific 16S gene regions. Samples which did not meet the sampling depth threshold of ≥ 1800 typically exhibited a markedly altered taxonomic composition, presumably because the low depth prevented most taxa from being sequenced. This highlights the importance of selecting an appropriate sampling depth cut-off to ensure that the data best reflects the bacterial diversity of each sample.

Contamination was a major focus of this study given its well-documented influence in microbiome research^45,46^ and the increased risk associated with undergraduate students’ limited experience in aseptic technique. We were interested not only in whether students introduced contaminating bacteria at a higher rate, but also the ability for this contamination to be computationally identified and removed post hoc. The main way in which we sought to identify contamination was through the use of extraction blank controls. Although students generated 19 extraction blank controls, only 7 of these produced any sequencing reads, and the extraction blank control with the highest number of reads captured only 21 features. Neither of the extraction blank controls produced by the researcher generated any sequencing reads. The lack of sequencing success observed for these samples was likely due to the very low quantity of DNA present after the PCR and cleanup steps of the protocol. While this result is somewhat reassuring as it suggests that very little contaminating bacterial DNA was introduced by the students or researcher, the lack of resulting sequencing data prevented us from using tools such as decontam to identify which species in the dataset were contaminants. Our use of Nanopore sequencing technology may have made contamination identification more difficult as all other major identification tools (e.g. Deblur, DADA2, UNOISE) are designed for low error rate short-read sequencing and as such were not appropriate for our dataset.

In order to gain a level of understanding of contamination in the absence of these tools, we manually inspected the genera present within the extraction blank controls. Of the 26 genera in the extraction blank controls, 22 were also present within experimental samples. The 10 most abundant of these genera – *Massilia, Streptococcus, Planomicrobium, Enterococcus,* an unclassified genus in the *Planococcaceae* family, *Solibacillus, Romboutsia,* an unclassified genus in the *Bacilli* class, *Paeniclostridium,* and an unclassified genus in the *Longimicrobiaceae* family – have all been identified within human or animal gut microbiomes^35–38,47–51^, and, crucially, as members of the echidna gut microbiota^18,34^. This suggests that instead of being contaminants introduced by the students, these genera were present within the experimental samples and crossed over into the extraction blank controls at some point during the microbiome workflow. This is not uncommon for high microbial biomass samples such as scats^46^. Four genera - *Aggregatibacter, Veillonella, Haemophilus,* and *Neisseria* - were unique to the extraction blank controls. These species are typically associated with the oral or gut microbiome^52–54^, and were either extremely lowly abundant or absent from existing echidna gut microbiome datasets^18,34^. These taxa may represent genuine contamination - but considering their low abundance within the extraction blank controls and absence from experimental samples, they are not cause for concern. Our results suggest that students do not introduce contamination that would significantly alter bacterial composition or diversity; however, the failure to sequence the majority of the extraction blank controls, coupled with the lack of post hoc contamination tools available for Nanopore sequencing data, mean that the true level of contamination could not be accurately quantified.

Despite the drawbacks that we have previously discussed, the use of Oxford Nanopore sequencing technology presented several benefits in an educational setting. The portable nature of the MinION™ sequencer meant that samples could be sequenced in the practical laboratory. Furthermore, this device provides live metrics of sequencing success. The interactivity and instant feedback inherent to this system provided a highly engaging format for educating students on cutting-edge genetics technology. The ability of teaching staff to conduct sequencing in-house allowed for fast turnaround times; with appropriate course timetabling, students could have the opportunity to analyse their own data. Although the use of portable sequencing devices in educational contexts has only recently been made possible, the benefits for student engagement and learning have already been described^55,56^. Designers of microbiome-based authentic research experiences should carefully weigh these benefits with the aforementioned drawbacks when selecting a sequencing approach.

The design of our study introduced statistical limitations. The presence of several student replicates but only one researcher replicate per scat sample caused inherent asymmetry within the dataset. This prevented us from directly comparing student and researcher diversity metrics as it was not possible to calculate variance estimates for the researcher group. Additionally, the fact that the same scat sample was processed multiple times, and by different numbers of student groups, meant that we could not directly compare student and researcher averages or make use of purpose-built microbiome analysis tools such as ANCOM. Future studies should incorporate additional researchers and maintain consistent student group numbers across samples in order to more comprehensively evaluate students’ ability to characterise microbiota.

## Conclusion

Generation of research-grade microbiome data in the undergraduate classroom allows for processing of more samples while providing a real-world research experience for students. Our data suggest that undergraduate students can generate high-quality, accurate data for microbiome research. In addition to the compositional similarities between researcher- and student-generated microbiome datasets, we did not find evidence that students introduce contamination at a higher rate than researchers, though differences were seen for experimental efficiency and alpha diversity metrics. In light of these findings, we make the following three recommendations: first, that at least 3 replicates should be generated per sample in order to increase the likelihood of successful sequencing and to identify and mitigate potential random error introduced by students; second, that extraction blank controls should be sequenced wherever possible to gain an understanding of contaminating bacteria introduced by students as well as cross-contamination between experimental samples; and third, that when performing taxonomic and diversity analyses, a high sampling depth should be selected to ensure a true representation of the microbiome, even at the expense of sample numbers. Our research affirms the mutual benefits to both students and researchers of authentic microbiome research experiences in the undergraduate classroom and encourages the implementation of these programs into the future.

## Supporting information

Sample metadata

Practical manuals

## Acknowledgements

The authors would like to acknowledge the Genetics IIIB practical classes of 2023 and 2024 for their enthusiastic participation in our laboratory sessions. We thank the School of Biological Sciences Teaching Technical Support Team for their invaluable assistance in implementing these practicals. IW was supported by a University of Adelaide Research Scholarship.

## Data availability

All metagenomic sequencing data is available at NCBI BioProjects accession number PRJNA1378557. Processed data as well as statistical analysis and visualisation code is available at the following GitHub repository: https://github.com/isa-wilson/microbiome_pracs/.

## Works Cited

1. Vuong, H. E., Yano, J. M., Fung, T. C. C, Hsiao, E. Y. The Microbiome and Host Behavior. Annu. Rev. Neurosci. 40, 21–49 (2017).

2. Comizzoli, P., Power, M. L., Bornbusch, S. L. C, Muletz-Wolz, C. R. Interactions between reproductive biology and microbiomes in wild animal species. *Anim*. Microbiome 3, 87 (2021).

3. Bronzo, V. et al. The Role of Innate Immune Response and Microbiome in Resilience of Dairy Cattle to Disease: The Mastitis Model. Animals 10, 1397 (2020).

4. Jin Song, S., et al. Engineering the microbiome for animal health and conservation. Exp. Biol. Med. 244, 494–504 (2019).

5. Kerkhof, L. J. Is Oxford Nanopore sequencing ready for analyzing complex microbiomes? FEMS Microbiol. Ecol. 97, (2021).

6. Wang, J. T. H. Course-based undergraduate research experiences in molecular biosciences—patterns, trends, and faculty support. FEMS Microbiol. Lett. 364, fnx157 (2017).

7. Brownell, S. E. C and Kloser, M. J. Toward a conceptual framework for measuring the effectiveness of course-based undergraduate research experiences in undergraduate biology. *Stud*. High. Educ. 40, 525–544 (2015).

8. Ing, M., BurnetteIII, J. M., Azzam, T. C, Wessler, S. R. Participation in a Course-Based Undergraduate Research Experience Results in Higher Grades in the Companion Lecture Course. Educ. Res. 50, 205–214 (2021).

9. Bangera, G. C, Brownell, S. E. Course-Based Undergraduate Research Experiences Can Make Scientific Research More Inclusive. CBE—Life Sci. Educ. 13, 602–606 (2014).

10. Kerr, M. A. C, Yan, F. Incorporating Course-Based Undergraduate Research Experiences into Analytical Chemistry Laboratory Curricula. J. Chem. Educ. 93, 658–662 (2016).

11. Muth, T. R. C, Caplan, A. J. Microbiomes for All. Front. Microbiol. 11, (2020).

12. Wang, M. et al. Student-Driven Course-Based Undergraduate Research Experience (CUREs) Projects in Identifying Vaginal Microorganism Species Communities to Promote Scientific Literacy Skills. Front. Public Health 10, (2022).

13. Baker, S. S. et al. Students in a Course-Based Undergraduate Research Experience Course Discovered Dramatic Changes in the Bacterial Community Composition Between Summer and Winter Lake Samples. Front. Microbiol. 12, (2021).

14. Zelaya, A. J., Gerardo, N. M., Blumer, L. S. C, Beck, C. W. The Bean Beetle Microbiome Project: A Course-Based Undergraduate Research Experience in Microbiology. Front. Microbiol. 11, (2020).

15. Aho, E. L., Ogle, J. M. C, Finck, A. M. The Human Microbiome as a Focus of Antibiotic Discovery: Neisseria mucosa Displays Activity Against Neisseria gonorrhoeae. Front. Microbiol. 11, (2020).

16. Sun, E. et al. Development of a data science CURE in microbiology using publicly available microbiome datasets. Front. Microbiol. 13, (2022).

17. Perry, T., et al. EchidnaCSI: Engaging the public in research and conservation of the short-beaked echidna. Proc. Natl. Acad. Sci. 119, e2108826119 (2022).

18. Perry, T. et al. Characterising the Gut Microbiomes in Wild and Captive Short-Beaked Echidnas Reveals Diet-Associated Changes. Front. Microbiol. 13, (2022).

19. Matoute, A. et al. Meat-Borne-Parasite: A Nanopore-Based Meta-Barcoding Work-Flow for Parasitic Microbiodiversity Assessment in the Wild Fauna of French Guiana. Curr. Issues Mol. Biol. 46, 3810–3821 (2024).

20. Bolyen, E. et al. Reproducible, interactive, scalable and extensible microbiome data science using QIIME 2. Nat. Biotechnol. 37, 852–857 (2019).

21. SILVA ribosomal RNA gene database project: improved data processing and web-based tools | Nucleic Acids Research | Oxford Academic. https://academic.oup.com/nar/article/41/D1/D590/1069277.

22. Wickham, H. Ggplot2: Elegant Graphics for Data Analysis. (Springer international publishing, Cham, 2016).

23. de Vries, A. C, Ripley, B. D. ggdendro: Create Dendrograms and Tree Diagrams Using ‘ggplot2’. (2024).

24. McMurdie, P. J. C, Holmes, S. phyloseq: An R Package for Reproducible Interactive Analysis and Graphics of Microbiome Census Data. PLOS ONE 8, e61217 (2013).

25. Barnett, D. J. m, Arts, I. C. w C, Penders, J. microViz: an R package for microbiome data visualization and statistics. J. Open Source Softw. 6, 3201 (2021).

26. Smith, S. schuyler-smith/phylosmith: Initial Release. Zenodo 10.5281/ZENODO.3251024 (2019).

27. Davis, N. M., Proctor, D. M., Holmes, S. P., Relman, D. A. C, Callahan, B. J. Simple statistical identification and removal of contaminant sequences in marker-gene and metagenomics data. Microbiome 6, 226 (2018).

28. Schrader, C., Schielke, A., Ellerbroek, L. C, Johne, R. PCR inhibitors – occurrence, properties and removal. J. Appl. Microbiol. 113, 1014–1026 (2012).

29. Eriksson, P., Mourkas, E., González-Acuna, D., Olsen, B. C, Ellström, P. Evaluation and optimization of microbial DNA extraction from fecal samples of wild Antarctic bird species. Infect. Ecol. Epidemiol. 7, 1386536 (2017).

30. Hyman*, O., et al. CURE-all: Large Scale Implementation of Authentic DNA Barcoding Research into First-Year Biology Curriculum. 10.24918/cs.2019.10 (2021) doi:10.24918/cs.2019.10.

31. Teng, F. et al. Impact of DNA extraction method and targeted 16S-rRNA hypervariable region on oral microbiota profiling. Sci. Rep. 8, 16321 (2018).

32. Starke, I. C., Vahjen, W., Pieper, R. C, Zentek, J. The Influence of DNA Extraction Procedure and Primer Set on the Bacterial Community Analysis by Pyrosequencing of Barcoded 16S rRNA Gene Amplicons. Mol. Biol. Int. 2014, 548683 (2014).

33. Luna, G. M., Dell’Anno, A. C, Danovaro, R. DNA extraction procedure: a critical issue for bacterial diversity assessment in marine sediments. Environ. Microbiol. 8, 308– 320 (2006).

34. Buthgamuwa, I. et al. Gut microbiota in the short-beaked echidna (Tachyglossus Aculeatus) shows stability across gestation. MicrobiologyOpen 12, e1392 (2023).

35. Piquer-Esteban, S., Ruiz-Ruiz, S., Arnau, V., Diaz, W. C, Moya, A. Exploring the universal healthy human gut microbiota around the World. Comput. Struct. Biotechnol. J. 20, 421–433 (2022).

36. Krawczyk, B., Wityk, P., Gałęcka, M. C, Michalik, M. The Many Faces of Enterococcus spp.—Commensal, Probiotic and Opportunistic Pathogen. Microorganisms 9, 1900 (2021).

37. Rojas, C. A. et al. Taxonomic, Genomic, and Functional Variation in the Gut Microbiomes of Wild Spotted Hyenas Across 2 Decades of Study. mSystems 8, e00965–22 (2022).

38. Xi, L. et al. Comparative study of the gut microbiota in three captive Rhinopithecus species. BMC Genomics 24, 398 (2023).

39. Zhang, M.-L., et al. Citrobacter Species Increase Energy Harvest by Modulating Intestinal Microbiota in Fish: Nondominant Species Play Important Functions. mSystems 5, 10.1128/msystems.00303-20 (2020).

40. Hamilton, A. L., Kamm, M. A., Ng, S. C. C, Morrison, M. Proteus spp. as Putative Gastrointestinal Pathogens. Clin. Microbiol. Rev. 31, 10.1128/cmr.00085-17 (2018).

41. Pasolli, E. et al. Large-scale genome-wide analysis links lactic acid bacteria from food with the gut microbiome. Nat. Commun. 11, 2610 (2020).

42. Hoyles, L. et al. Recognition of greater diversity of *Bacillus* species and related bacteria in human faeces. Res. Microbiol. 163, 3–13 (2012).

43. Zhang, Z. et al. A Diversified Dietary Pattern Is Associated With a Balanced Gut Microbial Composition of Faecalibacterium and Escherichia/Shigella in Patients With Crohn’s Disease in Remission. J. Crohns Colitis 14, 1547–1557 (2020).

44. Jabeen, I., Islam, S., Hassan, A. K. M. I., Tasnim, Z. C, Shuvo, S. R. A brief insight into Citrobacter species - a growing threat to public health. Front. Antibiot. 2, 1276982 (2023).

45. Lauder, A. P. et al. Comparison of placenta samples with contamination controls does not provide evidence for a distinct placenta microbiota. Microbiome 4, 29 (2016).

46. Eisenhofer, R. et al. Contamination in Low Microbial Biomass Microbiome Studies: Issues and Recommendations. Trends Microbiol. 27, 105–117 (2019).

47. Zhou, T., Liu, S. C, Jiang, A. Comparison of gut microbiota between immigrant and native populations of the Silver-eared Mesia (Leiothrix argentauris) living in mining area. Front. Microbiol. 14, (2023).

48. Qi, W., et al. Integrated Analysis of the Transcriptome and Microbial Diversity in the Intestine of Miniature Pig Obesity Model. Microorganisms 12, 369 (2024).

49. Magruder, M. et al. Gut commensal microbiota and decreased risk for Enterobacteriaceae bacteriuria and urinary tract infection. Gut Microbes 12, 1805281 (2020).

50. Rothenberg, S. E. et al. Fecal Methylmercury Correlates With Gut Microbiota Taxa in Pacific Walruses (Odobenus rosmarus divergens). Front. Microbiol. 12, (2021).

51. Giraud, C. et al. The Active Microbiota of the Eggs and the Nauplii of the Pacific Blue Shrimp Litopenaeus stylirostris Partially Shaped by a Potential Vertical Transmission. Front. Microbiol. 13, (2022).

52. Giacomini, J. J. et al. Spatial ecology of Haemophilus and Aggregatibacter in the human oral cavity. Microbiol. Spectr. 12, e04017–23 (2024).

53. Kolenbrander, P. The Genus Veillonella. in The Prokaryotes: Volume 4: Bacteria: Firmicutes, Cyanobacteria (eds Dworkin, M., Falkow, S., Rosenberg, E., Schleifer, K. - H. C, Stackebrandt, E.) 1022–1040 (Springer US, New York, NY, 2006). doi:10.1007/0-387-30744-3_36.

54. Van Dijck, C., Laumen, J. G. E., Manoharan-Basil, S. S. C, Kenyon, C. Commensal Neisseria Are Shared between Sexual Partners: Implications for Gonococcal and Meningococcal Antimicrobial Resistance. Pathogens 9, 228 (2020).

55. Callura, J. Bridging Biology and Data Science: Nanopore Sequencing-Based Lab Module for Biocomputational Engineering Students. *Biomed*. Eng. Educ. 10.1007/s43683-025-00196-4 (2025) doi:10.1007/s43683-025-00196-4.

56. Cervantes, J., Perry, C. C, Wang, M. C. Teaching next-generation sequencing to medical students with aCnbsp;portable sequencing device. Perspect. Med. Educ. 10, 252–255 (2020).

