## Supplementary Figures for "Undergraduate student practicals generate high-quality data for microbiome research"

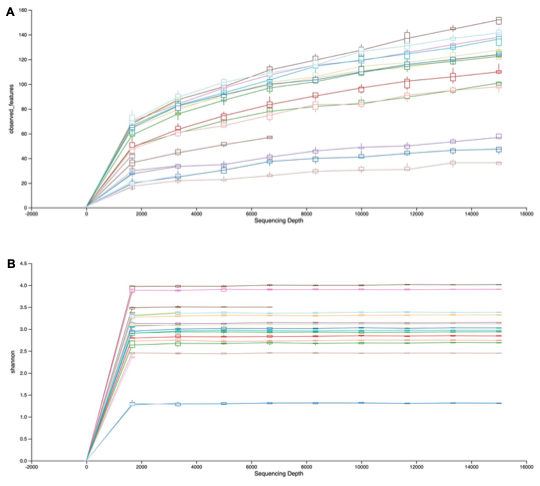

**Figure S1:** Alpha rarefaction curves showing **(A)** observed OTUs and **(B)** Shannon diversity at various sequencing depths. Each line represents one sample.

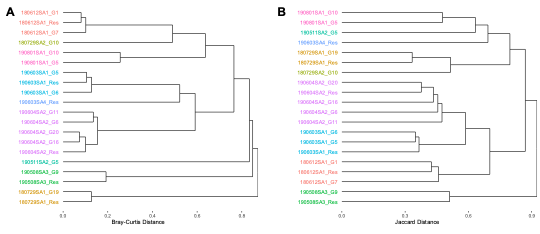
**Figure S2:** Dendrograms depicting hierarchical clustering of sample replicates using **(A)** Bray-Curtis distances and **(B)** Jaccard distances. Replicates are coloured according to scat ID.
