## Supplementary material for "Undergraduate student practicals generate high-quality data for microbiome research": Practical manuals: PracManual_Session1.pdf

### Session 1: DNA extraction from echidna scats

In this practical session you will extract bacterial DNA from wild echidna scats collected in the Flinders Ranges through the EchidnaCSI citizen science project. You will then assess the quality of your DNA extractions using gel electrophoresis.

**Materials (per 2 student pair):**

- 2 x esky containing dry ice
- 4 x mortars and pestles (stored in fridge/freezer until ready to use)
- 4 x metal spatulas
- Eppendorf tubes – 1.5 mL and 2 mL
- Pipettes – p1000 and p200
- **Filter** pipette tips – 1000 μL and 200 μL
- 5 x Spin Columns in collection tube (keep in packaging until ready to use)
- Spare collection tubes
- Reagents – InhibitEX buffer, Proteinase K, Buffer AL, Ethanol (100%), Buffer AW1, Buffer AW2, Buffer ATE
- Textas

!

**Prepare your workspace**

*It is VERY important that your bench and equipment are clean when working with microbial DNA in order to avoid contamination.*

Use 70% ethanol to **wipe down** your bench, pipettes, pipette tip boxes, tube boxes, racks, pens/textas, etc – anything that you will be touching while carrying out the experiments. Ensure that you are **using gloves at all times** to minimise contamination from your skin or other sources.

**Prepare your samples**

*Each student will be provided with two scat samples to extract DNA from. Each sample has a unique ID number. Each pair will also prepare one Extraction Blank Control (EBC) to which you will not add a sample.*

1. On the piece of paper provided, write down your university ID (‘a’ number).
2. Write down the four ID numbers of your scat samples.
3. On the space labelled “Extraction Blank Control”, write an ID number that is “EBC” followed by a combination of you and your partner’s initials (e.g. EBC-IWRV).

**Prepare your tubes**

*Each pair should prepare ten 2 mL tubes: two for each sample plus one for the EBC.*

1. Label the ten tubes as follows:
    - Five tubes: InhibitEX + sample ID or EBC ID
    - Five tubes: ProtK + sample ID or EBC ID
2. Pipette **1.4 mL InhibitEX** buffer into each 2 mL tube labelled with "InhibitEX" (=5 tubes).
3. Pipette **25 µL Proteinase K** into each 2 mL tube labelled with "ProtK" (=5 tubes).

**Homogenise the samples**

*Each student will repeat this process twice, using a* ***fresh mortar and pestle*** *for each of the two scat samples. Each pair will homogenise four scats in total*.

1. Nestle a clean mortar and pestle into an esky filled with dry ice.
2. Pour the sample into the mortar and pestle and grind for around 5-10 minutes until homogenised.
    - Tick on the checklist which sample you have just placed into the mortar and pestle.
    - Using the pestle, first tamp (press firmly) on the sample to break it up then use circular motions to crush. It is helpful to occasionally use the side of the mortar/pestle.
    - Indicators that you are done:
        - There are no visible sparkly pieces
        - The sample will be visibly lighter
        - It will be the consistency of flour or icing sugar: it will not feel gritty when grinding
    - Once you think you are done, keep going for another two minutes.
3. Using a clean metal spatula, scoop the sample into the prepared "InhibitEX + sample ID" tube until the tube is approximately 1/3 full of sample.
    - Flick and invert the tube a few times to ensure that the sample is completely covered by the buffer (no dry spots).
    - **Change gloves before moving on to your next sample**.
4. Repeat steps 1-3 for your next sample.
5. For all five "InhibitEX + ID) tubes (including the EBC), vortex for 1 min then centrifuge for 3 min. *Note: all centrifugation steps in this protocol are carried out at max speed.*

**Extract the DNA**

1. Label four 1.5 mL tubes with the four scat sample IDs.
2. Transfer **800 µL supernatant** into the 1.5 mL tubes, trying not to pick up any solid material. Discard the original 2 mL tube. Centrifuge for 3 min.
3. Transfer **600 µL supernatant** from the 1.5 mL tubes into the prepared 2 mL "ProtK + sample ID" tubes. This time, be very careful not to pick up any solid material or floating particles (it is better to pipette up less liquid than to carry over solid material).
4. Transfer over **600 µL** of the "InhibitEX + EBC ID" sample to the "ProtK + EBC ID" tube.

*From here onwards, carry out the following steps the same with all of your tubes – your four scat samples and your EBC*.

5. Add **600 µL Buffer AL** and vortex for 15 s until homogenous. *Do not add Buffer AL directly to proteinase K.*
6. Incubate at 70°C for 10 min.
7. Add **600 µL ethanol (100%)** and mix by vortexing.
8. Label five spin columns with your IDs (4 x scat samples + 1 EBC).
9. Carefully add **600 µL lysate** from the "ProtK + ID" tubes into the labelled spin columns. Close the cap and centrifuge for 1 min. Place the spin column into a new collection tube. Discard the old collection tube containing the filtrate.
10. Repeat the previous step twice more until all of the lysate has passed through the spin column. Place the spin column into a new collection tube. Discard the old collection tube containing the filtrate.
11. Carefully open the spin column and add **500 µL Buffer AW1**. Centrifuge for 1 min. Place the spin column into a new collection tube. Discard the old collection tube containing the filtrate.
12. Add **500 µL Buffer AW2** to the spin column. Centrifuge for 3 min. Place the spin column into a new collection tube. Discard the old collection tube containing the filtrate.
13. Centrifuge for 3 min. *This step helps to eliminate any residual buffer from the column*.
14. Label five 1.5 mL tubes with your IDs (4x scat samples + 1 EBC). *This time make sure to label the top and side of the tube. Add your name to the side of the tube too. The labels must be very clear/legible. This is where your extracted DNA will be stored*.
15. Place the spin columns into the newly labelled 1.5 mL tubes. Pipette **200 µL Buffer ATE** directly onto the membrane of the spin column. Incubate for 1 min at room temperature. Centrifuge for 1 min to elute DNA.

**Assess DNA quality using gel electrophoresis**

1. Add **9 µL of your extracted DNA** and **1 µL of 10X loading buffer** to a 1.5 mL tube. *Do this for each of your samples including the EBC*.
2. Mix gently by pipetting up and down.
3. Load the whole sample onto the gel. *Write down the sample ID and lane number on the sheet provided next to the gel tanks*.
4. Run for 30 min at 100 V.
