## Supplementary material for "Undergraduate student practicals generate high-quality data for microbiome research": Practical manuals: PracManual_Session2.pdf

### Protocol: Library prep for Nanopore sequencing

In the previous session you extracted DNA from echidna scat samples. Before this session the demonstrators performed PCR using your extracted DNA to amplify the 16S gene from bacterial DNA present within your samples.

In this session you will be preparing the 16S amplicons for sequencing on a Nanopore MinION. You will first quantify your 16S amplicons samples using a Qubit Fluorometer. The demonstrators will use these concentrations to pool your samples together. The pools will be cleaned using magnetic beads, which removes any contaminants that could interfere with sequencing. The demonstrators will add sequencing reagents to the pools then begin sequencing on a Nanopore MinION. *Please note that while we may not be able to sequence everyone's samples today, we will sequence them over the next week or so and provide the data via MyUni*.

#### 16S Amplicon quantification (Qubit)

**Materials**

- 16S PCR products from your samples
- Qubit assay tubes
- Qubit working solution (WS)
- **FILTER** pipette tips (p200)

You will prepare one Qubit assay tube for each of your cleaned samples.

1. Label the lid of a Qubit assay tube with your sample ID. ***Do not label the side of the tube***. *Labelling the side of the tube may interfere with quantification.*

2. Add **199 µL** of the **Qubit working solution (WS)** to each **Qubit assay tube**. Place the Qubit tubes in a drawer - they should be kept away from light until they are ready for quantification.

3. The demonstrators will call up one pair at a time to bring their samples over to the Qubit fluorometer for quantification. Bring your prepared Qubit tubes and your cleaned samples.

4. Add 1 µL of your cleaned sample to its Qubit tube. Vortex for 3-5 seconds. Be careful not to create bubbles. Ensure that there are no droplets on the sides or lid of the tube.

5. Incubate the tubes for 2 minutes at room temperature then quantify using the Qubit fluorometer.

#### Sample pooling & magnetic bead clean-up

*Please observe the demonstrators as they carry out these steps.*

The DNA quantifications from the previous step will be used to combine your samples at an equimolar concentration in preparation for sequencing. These samples will then be “cleaned” using magnetic beads. The demonstrators will measure the concentration of the cleaned DNA pools using the Qubit.

#### Library preparation

The demonstrators will add sequencing reagents to the pools before loading them onto a Nanopore flow cell for sequencing.
