## Supplementary material for "Undergraduate student practicals generate high-quality data for microbiome research": Practical manuals: Session1_Checklist.pdf

### DNA EXTRACTION

### CHECKLIST

Name: ______________________ Uni ID: ______________

Name: ______________________ Uni ID: ______________

FILL IN THE CHECKLIST BELOW WITH EVERY TASK YOU COMPLETE AS YOU MOVE THROUGH THE PROTOCOL.

| | Student 1 | | Student 2 | | |
|---|---|---|---|---|---|
| | Scat ID#1: ______ | Scat ID#2: ______ | Scat ID#3: ______ | Scat ID#4: ______ | EBC ID: ______ |
| • 2mL tube labelled 'InhibitEX + ID' | ☐ | ☐ | ☐ | ☐ | ☐ |
| • 1.4mL InhibitEX buffer added to above tube | ☐ | ☐ | ☐ | ☐ | ☐ |
| • 2mL tube labelled 'ProtK + ID' | ☐ | ☐ | ☐ | ☐ | ☐ |
| • 25uL ProtK added to above tube | ☐ | ☐ | ☐ | ☐ | ☐ |
| • Scat placed in mortar | ☐ | ☐ | ☐ | ☐ | |
| • Scat scooped into tube 'InhibitEX + ID' | ☐ | ☐ | ☐ | ☐ | |
| (*only fill one third of tube with scat*) | | | | | |
| • Tube inverted to mix fully with buffer | ☐ | ☐ | ☐ | ☐ | |
| • Vortexed for 1 min | ☐ | ☐ | ☐ | ☐ | ☐ |
| • Centrifuged for 3 mins | ☐ | ☐ | ☐ | ☐ | ☐ |
| • 1.5mL tube labelled with sample ID | ☐ | ☐ | ☐ | ☐ | |
| • 800uL eluate transferred to above tube | ☐ | ☐ | ☐ | ☐ | |
| • Centrifuged for 3 mins | ☐ | ☐ | ☐ | ☐ | |
| • 600uL eluate transferred to tube 'ProtK + ID' | ☐ | ☐ | ☐ | ☐ | ☐ |
| (*DO NOT transfer solid particles over*) | | | | | |
| • 600uL of Buffer AL added | ☐ | ☐ | ☐ | ☐ | ☐ |
| • Vortexed for 15s | ☐ | ☐ | ☐ | ☐ | ☐ |
| • Placed at 70 degrees for 10 mins | ☐ | ☐ | ☐ | ☐ | ☐ |
| • 600uL ethanol added to tube & vortexed | ☐ | ☐ | ☐ | ☐ | ☐ |

### DNA EXTRACTION

### CHECKLIST

| | Student 1 | | Student 2 | | |
|---|---|---|---|---|---|
| | Scat ID#1: | Scat ID#2: | Scat ID#3: | Scat ID#4: | EBC ID: |
| • Spin column labelled with 'ID' | ☐ | ☐ | ☐ | ☐ | ☐ |
| • 600uL of eluate added to spin column | ☐ | ☐ | ☐ | ☐ | ☐ |
| • Centrifuged for 1 min | ☐ | ☐ | ☐ | ☐ | ☐ |
| • Spin column moved to new collection tube | ☐ | ☐ | ☐ | ☐ | ☐ |
| • Above 3 steps repeated once | ☐ | ☐ | ☐ | ☐ | ☐ |
| • Above 3 steps repeated twice | ☐ | ☐ | ☐ | ☐ | ☐ |
| • Spin column moved to new collection tube | ☐ | ☐ | ☐ | ☐ | ☐ |
| • 500uL Buffer AW1 added | ☐ | ☐ | ☐ | ☐ | ☐ |
| • Centrifuged for 1 min | ☐ | ☐ | ☐ | ☐ | ☐ |
| • Spin column moved to new collection tube | ☐ | ☐ | ☐ | ☐ | ☐ |
| • 500uL Buffer AW2 added | ☐ | ☐ | ☐ | ☐ | ☐ |
| • Centrifuged for 3 mins | ☐ | ☐ | ☐ | ☐ | ☐ |
| • Spin column moved to new collection tube | ☐ | ☐ | ☐ | ☐ | ☐ |
| • Centrifuged for 3 mins (nothing added) | ☐ | ☐ | ☐ | ☐ | ☐ |
| • 1.5mL tube labelled with 'ID' + your name (*Write on lid & side of tube and make sure handwriting is clear*) | ☐ | ☐ | ☐ | ☐ | ☐ |
| • Spin column added to labelled 1.5mL tube | ☐ | ☐ | ☐ | ☐ | ☐ |
| • 200uL of Buffer ATE added | ☐ | ☐ | ☐ | ☐ | ☐ |
| • Incubated at room temp for 1 min | ☐ | ☐ | ☐ | ☐ | ☐ |
| • Centrifuged for 1 min | ☐ | ☐ | ☐ | ☐ | ☐ |
| • Keep 1.5mL tube & discard spin column | ☐ | ☐ | ☐ | ☐ | ☐ |
